# Field-evolved resistance to neonicotinoids in the mosquito, *Anopheles gambiae*, is associated with downregulation and mutations of nicotinic acetylcholine receptor subunits combined with cytochrome P450-mediated detoxification

**DOI:** 10.1101/2024.08.17.608399

**Authors:** Caroline Fouet, Matthew J. Pinch, Fred A. Ashu, Marilene M. Ambadiang, Calmes Bouaka, Anthoni J. Batronie, Cesar A. Hernandez, Desiree E. Rios, Véronique Penlap-Beng, Colince Kamdem

## Abstract

Neonicotinoid insecticides act selectively on their nicotinic receptor targets leading to variable sensitivity among arthropods. This study aimed to investigate the molecular mechanisms underlying contrasting susceptibility to neonicotinoids observed in wild populations of two mosquito sibling species. Bioassays and a synergism test revealed that the sister taxa, *Anopheles gambiae* and *An. coluzzii*, from Yaounde, Cameroon, rely on cytochrome P450s to detoxify neonicotinoids and develop resistance. However, contrary to *An. coluzzii*, *An. gambiae* populations are evolving stronger resistance to several active ingredients facilitated by mutations and reduced expression of nicotinic acetylcholine receptors. Six mutations were detected in coding sequences of the β1 and α6 subunits, including two substitutions in one of the loops that modulate ligand binding and sensitivity. Allele frequencies were strongly correlated with a susceptibility gradient between *An. coluzzii* and *An. gambiae* suggesting that the mutations may play a key role in sensitivity. Messenger RNA expression levels of the β1, α3, and α7 subunits decreased dramatically, on average by 23.27, 17.50, 15.80-fold, respectively, in wild *An. gambiae* populations compared to a susceptible insectary colony. By contrast, only the β2 and α9-1 subunits were moderately downregulated (5.28 and 2.67-fold change, respectively) in field-collected *An. coluzzii* adults relative to susceptible colonized mosquitoes. Our findings provide critical information for the application and resistance management of neonicotinoids in malaria prevention.

## 1 Introduction

The malaria mosquito *Anopheles gambiae* is a species complex containing the most efficient vectors of *Plasmodium* in sub-Saharan Africa [1]. This complex includes at least nine cryptic taxa with varying vectorial capacities that have spread in different niches throughout the continent [2–4]. *Anopheles gambiae* sensu stricto (hereafter referred to as *Anopheles gambiae*) and *An. coluzzii* are two young species of the complex exhibiting exceptional adaptation to human habitats and vector control strategies [2,5–7]. Neurotoxic insecticides have been widely used over the past two decades in malaria prevention programs involving the deployment of insecticidal treated nets and indoor residual spraying of insecticides in sub-Saharan Africa [8]. Mosquito populations have rapidly developed adaptive strategies to cope with persistent exposure to lethal doses of synthetic chemicals [9,10]. Some classes of insecticides are simultaneously used in agriculture and in public health programs leading to cross-resistance [11]. The major modes of adaptation underlying resistance in *Anopheles* mosquitoes include modifications of the target site which reduce its affinity for the toxicant and overexpression of enzymes that metabolize and detoxify the insecticide [12,13].

There is significant heterogeneity in the intensity of phenotypic resistance and its molecular determinants among mosquito vector species [10]. For example, mutations of the voltage-gated sodium channel involved in resistance to pyrethroids and DDT have spread in *An. gambiae* for several decades, but were only recently discovered in one population of *An. funestus*, another major vector with near continent-wide distribution [14,15]. Even within species, local populations differ in how they use metabolic detoxification, target-site mutations or both mechanisms to adapt to a given insecticide [16–18]. Geographic variations in the intensity of selective pressures play a prime role in the development of such heterogeneity. Anopheline species are distributed across extremely diverse environmental settings where larval and adult populations are exposed to different insecticide selection pressures, which determine their ability to develop phenotypic and molecular resistance [19–22]. A better understanding of the mechanisms underlying the intensiity of resistance in vector species is key to effective deployment of insecticide-based malaria prevention tools.

Neonicotinoids are a class of eight insecticides applied in agriculture worldwide for at least three decades [23,24]. Formulations of clothianidin or imidacloprid, two neonicotinoids used in agricultural pest management, have recently been prequalified for malaria prevention interventions [25]. Evaluation of baseline susceptibility to neonicotinoids such as clothianidin, imidacloprid, acetamiprid and thiamethoxam has indicated a substantial variation in sensitivity among wild anopheline populations [26–32]. Residual exposure to neonicotinoids used for crop protection in agricultural areas is suspected to be a driver of cross-resistance to several active ingredients in larvae and adults [28]. Some wild populations of another major vector, *An. funestus*, also show signals of reduced susceptibility to acetamiprid, imidacloprid and clothianidin although larvae of this species occur in permanent breeding sites that are less exposed to residual pesticides [30]. This suggests that in addition to varying selection pressures, neonicotinoids may have differential toxicity towards *Anopheles* species, and that cross-resistance with other classes of insecticides could also be a contributing factor. Studying closely related taxa in small geographic areas allowing us to monitor the potential environmental pressures could provide insights into the drivers and the molecular changes underlying differential susceptibility to neonicotinoids in malaria vectors. The sibling species *An. gambiae* and *An. coluzzii* are the most important vectors of *Plasmodium* in human-dominated environments in Africa. In Yaounde, Cameroon, populations of both species display contrasting susceptibility to neonicotinoid insecticides [27,29]. Contrary to *An. coluzzii*, *An. gambiae* larvae from rural areas of Yaounde can grow and emerge in water containing a dose of acetamiprid, imidacloprid or clothianidin that kills immature stages of a susceptible insectary colony within 24 h of exposure [28]. Based on the geographic distribution of the sister taxa, the candidate environmental drivers of this divergence are urbanization and agriculture, but the molecular mechanisms remain unexplored.

Resistance to neonicotinoids has been studied in diverse insect species [23,33]. The most common resistance mechanisms include reduced sensitivity of the target site due to mutations and altered expression of nicotinic acetylcholine receptor subunits (nAChRs) as well as metabolic detoxification involving cytochrome P450s (CYPs) [23]. Neonicotinoids act as competitive modulators targeting ten to twelve α and non-α nAChR subunits present in insects [34]. Mutations known to affect the binding affinity of neonicotinoids and to contribute to resistance development have been identified on the β1, β2, α1, α3 and α6 subunits in insects including major agricultural pests [35–42]. In addition to amino acid changes, reduced expression of some subunits has been associated with increased tolerance to imidacloprid or to thiamethoxam for instance [41,43–46]. CYPs play a critical role in the development of resistance to neonicotinoid in insects, which has been demonstrated through synergism tests, in vitro metabolism of active ingredients, transcriptome analysis and gene knockdown [47–55]. Target-site insensitivity and metabolic resistance often co-occur in insects that have evolved high resistance to neonicotinoids, but the timing of these changes and their relative contributions remain poorly understood. Field-evolved resistance to acetamiprid, imidacloprid or clothianidin in *Anopheles* mosquitoes is only partially overcome through inhibition of CYPs with piperonyl butoxide (PBO), which suggests that reduced sensitivity of nAChRs could play a role in resistance development [27,29].

In this study, we quantified messenger RNA expression levels of eight nAChR subunits and screened mutations across coding sequences of β1 and α6 in *An. coluzzii* and *An. gambiae* populations exhibiting contrasting susceptibility to neonicotinoids. The synergistic effect of PBO suggests that susceptibility to acetamiprid can be restored in most populations through inhibition of CYPs whereas resistance to clothianidin involved a more complex mechanism. We detected mutations in Agβ1 and Agα6 whose allele frequencies were correlated with a resistance gradient including two substitutions that likely affect the binding affinity of neonicotinoids. We found that the decreasing susceptibility of *An. gambiae* is also facilitated by reduced expression of three nAChR subunits. The findings are highly significant for the application of neonicotinoids in mosquito control.

## 2 Materials and methods

### 2-1 Mosquito populations

Mosquitoes were sampled along an urban-to-rural gradient in Yaounde, Cameroon. Yaounde is a city founded in the 1888 within the equatorial forest in Central Africa. The climate is tropical with two rainy seasons and two dry seasons. The two cryptic species, *An. gambiae* and *An. coluzzii*, segregate along an urban-to-rural gradient in Yaounde. *An. coluzzii* is adapted to urban environments where immature stages are often found in polluted breeding sites in densely populated areas [5]. The sister taxon, *An. gambiae*, occurs in rural areas where larval populations are frequently exposed to residual pesticide due to intensive agricultural activities. Both species are sympatric in suburban areas.

Mosquito larvae were collected from four locations including two sites harboring exclusively *An. coluzzii* and two villages where only *An. gambiae* was present. *Anopheles coluzzii* mosquitoes were sampled from Etoa Meki (3°52’53’’ N, 11°31’40’’ E) and from Combattant (3°52’53’’ N, 11°31’40’’ E), two neighborhoods located ∼3 km from one another at the center of the city. *Anopheles gambiae* populations were obtained from Nkolondom (3°56’43’’ N, 11°3’01’’ E), a sub-urban neighborhood known for intensive cultivation of lettuce and aromatic herbs and from Lobo (3° 53ʹ 31ʺ N, 11° 13ʹ 49ʺ E), a typical rural area characterized by degraded forests. Larvae were collected from puddles in Lobo, Etoa Meki and Combatant and from standing water between furrows and ridges in a farm in Nkolondom using conventional sampling methods [56]. Immature stages were reared to adults in a controlled insectary environment, and 2-3 days old female adults were used for insecticide bioassays and molecular analyses. Mosquitoes from two insectary strains, *An. coluzzii* Ngousso and *An. gambiae* Kisumu, were also tested. Both strains colonized respectively in 2006 and 1975 are susceptible to insecticides used in malaria prevention interventions including neonicotinoids.

### 2-2 Insecticides and bioassays

The activity of two neonicotinoid insecticides, clothianidin and acetamiprid, was tested on females aged 2-3 days old using standard adult bioassays. Technical-grade formulations for both insecticides were purchased from Sigma-Aldrich, UK. Stock solutions were prepared using absolute ethanol and stored at +4°C. The susceptibility of *An. gambiae* and *An. coluzzii* mosquitoes was assessed using CDC bottle bioassays as previously described [27,29,57–60]. Previous studies have shown that a gradient of susceptibility to neonicotinoids can be detected among *An. gambiae* and *An. coluzzii* populations from Yaounde using 75 µg of active ingredient (AI)/bottle for acetamiprid and 150 µg of AI/bottle for clothianidin as discriminating concentrations [26,27,29,58]. Bottles were coated with 1 ml of the discriminating concentration of each insecticide following the CDC protocol [57]. After 1-h exposure of females to the insecticide, mortality was observed daily for 72 h. Bottles coated with 1 ml of absolute ethanol were used as a negative control. For each bioassay, mortality was calculated across four tests of 25 females each. Mortality in control tests were lower than 5% and were not corrected. Fisher’s exact test was used to assess variations in mortality rates across treatments.

## 2-3 Synergism test

To observe the effect of the cytochrome P450 inhibitor, PBO, on mortality, a synergism test was conducted. Bottles were first coated with PBO (4%), and female mosquitoes were exposed to the synergist for 1 h. Pre-exposed individuals were then tested with the insecticide to detect any rise in mortality due to the synergistic effect of PBO. Mosquitoes from Nkolondom were exposed to 75 µg of AI/bottle and 150 µg of AI/ bottle for acetamiprid and clothianidin respectively, which corresponded to the lethal concentrations (LC_99_) for the susceptible strain *An. gambiae* Kisumu [58]. Since *An. coluzzii* is more susceptible to neonicotinoids, LC_50_ (40 µg /bottle of clothianidin and 10 µg /bottle of acetamiprid) was used. Mortality was recorded 72 h post-exposure. All tests were performed with females aged 2-3 days that were collected as larvae and reared to adults in standard insectary conditions. A Fisher’s exact test was used to determine if the difference in mortality with or without pre-exposure to PBO was statistically significant.

### 2-4 DNA extraction and sequencing of Agβ1 and Agα6 subunits

Exons from two subunits, *Anopheles gambiae* β1 (Ag β1) and Agα6, were sequenced to detect mutations potentially involved in neonicotinoid sensitivity. Pairs of primers (Table 1) were designed based on the *An. gambiae* PEST reference sequence to target three exons on Agβ1 (exons 1, 2 and 3) and three on Agα6 (exons 7, 8 and 9). The Agβ1 primer pair encompassed coding sequences where five amino acid substitutions increasing tolerance to neonicotinoids have been detected in several insect pests [35,36,40,41]. The Agα6 sequence included the G275E substitution known to increase resistance to spinosad, another broad-spectrum insecticide targeting insect nAChRs [42,61]. Genomic DNA was extracted from individual female mosquitoes using the Qiagen DNeasy® Blood and Tissue kit (Qiagen, Germantown, MD, USA) following the manufacturer’s instructions. Species identification was conducted using a PCR method which discriminates *An. gambiae* and *An. coluzzii* based on a point mutation on the ribosomal DNA [62]. The PCR products of Agβ1 and Agα6 were purified and sequenced using forward and reverse primers.

**Table 1:**
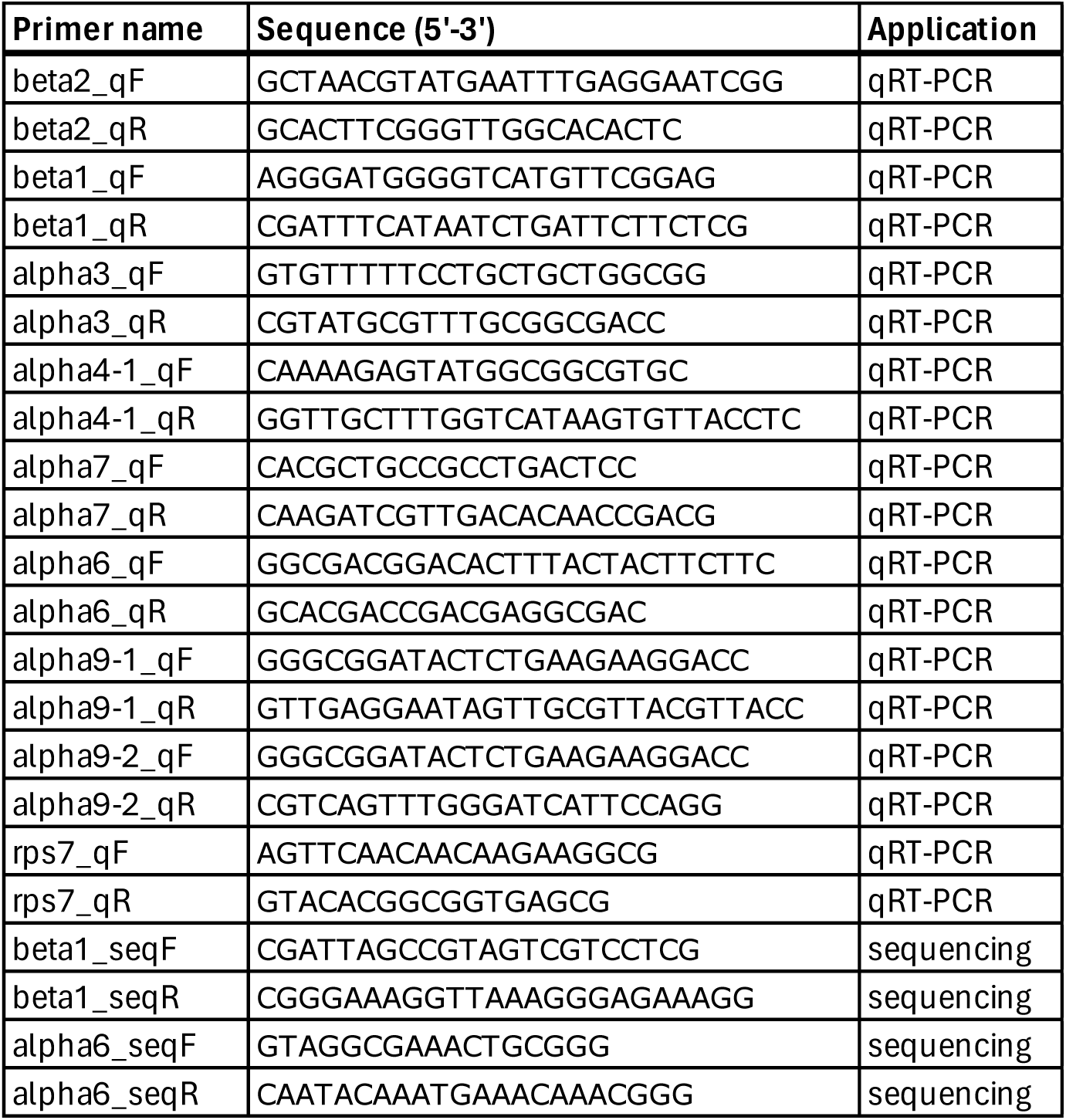
Primers used in this study.

Chromatograms were inspected and corrected. A consensus was created from the forward and reverse sequences using the program *Pearl* of the *Tracy* package. Multiple alignments were generated by aligning the consensus sequences to the *An. gambiae* PEST reference sequence using *Clustal W*, and nucleotide variations were observed in *Sabre*. Allele frequencies of mutations were compared between two susceptible laboratory strains, *An. gambiae* Kisumu and *An. coluzzii* Ngousso, non-exposed field-collected *An. gambiae* and *An. coluzzii* females, as well as wild *An. gambiae* mosquitoes alive 72 h post-exposure to acetamiprid or to clothianidin (LC_99_). Differences in allele frequencies between populations were assessed using a Fisher’s exact test.

### 2-5 RNA extraction and cDNA synthesis

For each population, RNA was extracted from pools of five 2-day-old female mosquitoes killed at the same time of day and stored in RNAlater (Invitrogen, Waltham, MA) at -80°C. Mosquitoes were homogenized in Buffer RLT (Qiagen, Germantown, MD, USA) containing beta-mercaptoethanol, and total RNA was isolated using the Qiagen RNeasy® kits (Qiagen, Germantown, MD, USA) according to the manufacturer’s protocol. All samples received an on-column treatment with RNA-free DNAse (Qiagen, Germantown, MD, USA) following the instructions in the RNeasy® kit. Isolated total RNA was quantified using the Qubit^TM^ RNA BR Assay kits (Thermo Fisher, Waltham, MA, USA) on a Qubit^TM^ 4 fluorometer (Thermo Fisher, Waltham, MA, USA). First-strand complementary DNA (cDNA) was synthesized from the total RNA using the iScript^TM^ Reverse Transcription Supermix (Bio-Rad, Hercules, CA, USA) following the manufacturer’s instructions. One hundred and fifty nanograms of total RNA was used as template.

### 2-6 mRNA expression levels of nAChR subunits

The mRNA expression levels of eight subunits (β1, β2, α3, α4-1, α6, α7, α9-1, α9-2) were measured using quantitative real-time polymerase chain reaction (qRT-PCR). Primer pairs (Table 1) were designed based on the *An. gambiae* PEST reference sequence. qRT-PCR reactions were done using iTaq™ Universal SYBR® Green (Bio-Rad, Hercules, CA, USA) with 20 μl reactions containing 1uL of cDNA, 100nM of each primer in a CFX 96 Opus Real Time System (Bio-Rad, Hercules, CA, USA). Initial denaturation was conducted at 95°C for 30 s, followed by 40 amplification cycles of denaturation at 95°C for 5 s and combined annealing and elongation at 60°C for 30 s. The fluorescence signal was measured at the end of each extension step at 60 °C. After amplification, a melting curve analysis was conducted to detect primer dimers. Samples were heated across a temperature gradient from 65 to 95°C at 0.5°C intervals, being held at each temperature for five seconds and then fluorescence signal was measured. PCR products were also verified using electrophoresis on agarose gel and sequencing. The gene, *ribosomal protein S7* (*rps7*), was used as internal reference gene for qPCR data normalization. The baseline mRNA expression levels for the eight subunits were compared between a susceptible colonized strain of *An. gambiae* (Kisumu) and two field populations of the same species (Lobo and Nkolondom). The levels of expression in *An. coluzzii* females from Etoa Meki and from Combattant were also compared to the baseline values detected in the susceptible strain Ngousso. Each population was represented by three biological replicates. The relative expression of each nAChR subunit was calculated based on the 2^−ΔΔC^ method [63]. The mRNA expression levels were analyzed with Welch t-tests using log-transformed values of fold-change. The level of significance was set at p < 0.05.

## 3 Results

### 3-1 *An. gambiae* is less susceptible to neonicotinoids than *An. coluzzii* in Yaounde

Mortality rate was evaluated in two susceptible strains and in four field populations exposed to 75 µg /ml of acetamiprid or to 150 µg /ml of clothianidin. Within 24 h of insecticide exposure, mortality reached 100% in *An. gambiae* Kisumu and *An. coluzzii* Ngousso consistent with the high susceptibility of both laboratory strains to acetamiprid and to clothianidin (Figure 1). Both *An. coluzzii* populations from urban areas of Yaounde (Etoa Meki and Combattant) were also highly susceptible to clothianidin reaching at least 98% mortality within 72 h of exposure. Mortality against acetamiprid at 72 h post-exposure was 86 ± 6.0 % in mosquitoes from Combattant and 88 ± 5.4 % in females from Etoa Meki. By contrast, both *An. gambiae* field populations showed low mortality to acetamiprid and to clothianidin indicating reduced susceptibility. Only 69.9 ± 9.5 % and 49.5 ± 6.6 % mortality were recorded within 72 h of exposure to clothianidin and acetamiprid respectively in females from Nkolondom. Mean mortality for acetamiprid was 87.0 ± 3.8 % in field-collected *An. coluzzii* versus only 58.0 ± 4.7 % in wild *An. gambiae* mosquitoes (Fisher’s exact test, p < 0.001, OR = 0.16, 95% CI = 0.096-0.260). Likewise, mean mortality for clothianidin was significantly lower (53 ± 7.4 %) in *An. gambiae* field populations compared to *An. coluzzii* mosquitoes from urban areas (99.3 ± 0.4 %) (Fisher’s exact test, p < 0.001, OR = 0.010, 95% CI = 0.001-0.036).

**Figure 1:**
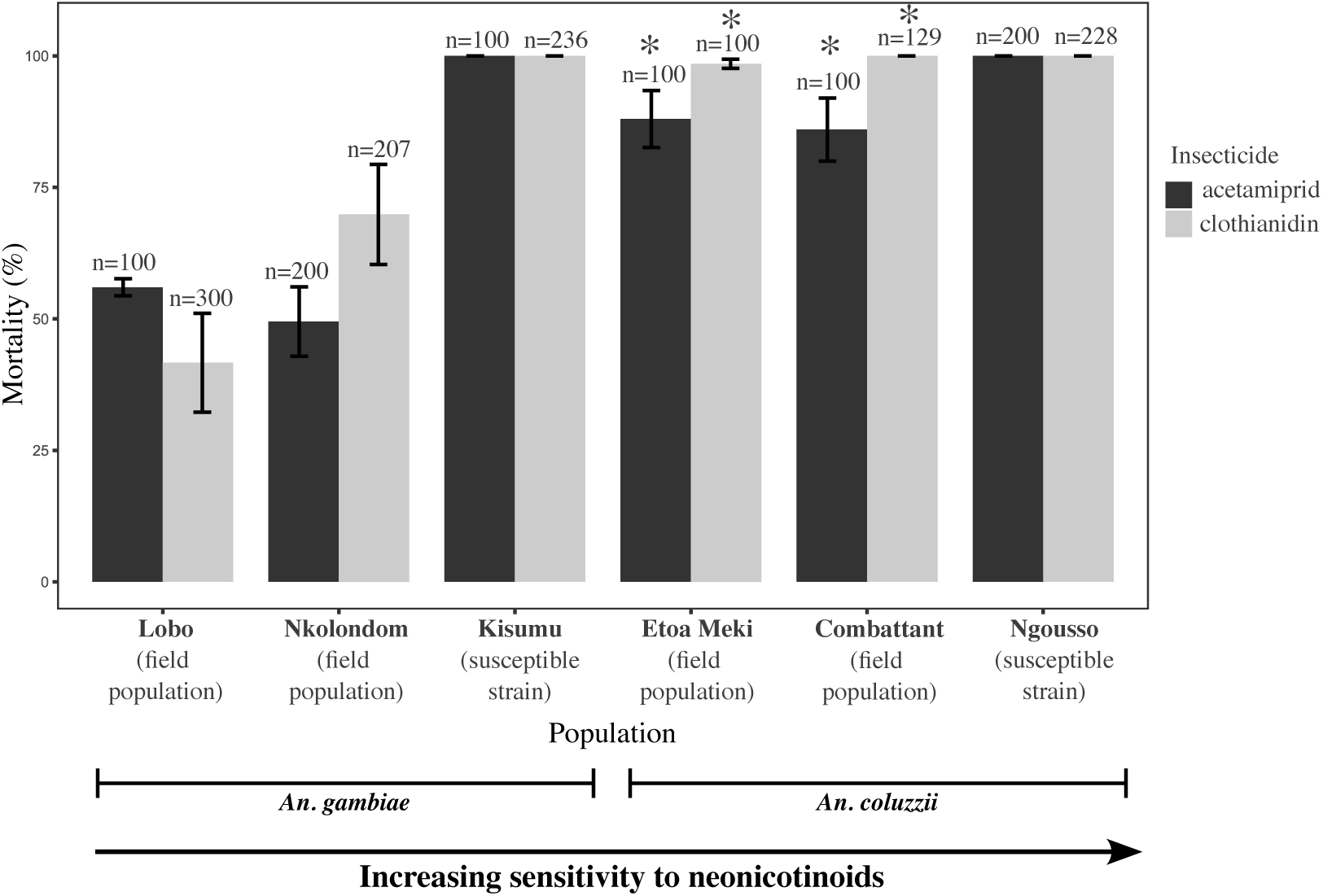
Mortality values reflecting a gradient of susceptibility to acetamiprid and clothianidin between *An. gambiae* and *An. coluzzii* populations. Mortality was recorded 72 h post-exposure to acetamiprid (75 µg /ml) or to clothianidin (150 µg /ml). Error bars represent the standard error of the mean. Asterisk denotes a significant difference in mortality between field-collected populations of *An. coluzzii* and *An. gambiae* (Fisher’s exact test, p < 0.05).

### 3-2 Piperonyl butoxide synergizes the activity of acetamiprid and clothianidin in *An. coluzzii* and *An. gambiae*

We performed a synergism test to determine if inhibiting cytochrome P450 enzymes with PBO can increase the efficacy of clothianidin and acetamiprid. Mortality rose from 49.5 ± 6.6% to 98.3 ± 1.8% (Fisher’s exact test, p < 0.001, OR = 56.1, 95% CI = 14.4-481) in *An. gambiae* field mosquitoes respectively exposed to acetamiprid (75 µg /ml) or pre-exposed to PBO before being tested with the active ingredient (Figure 2). Pre-exposure to PBO also increased mortality to clothianidin (150 µg /ml) in field-collected *An. gambiae* females (69.9 ± 9.5 % vs 87.5 ± 4.8%, Fisher’s exact test, p < 0.001, OR = 3.0, 95% CI = 1.50-6.51). The synergistic effect of PBO was also verified in susceptible mosquitoes using LC_50_ (40 µg /ml of clothianidin and 15 µg /ml of acetamiprid). Results showed that PBO enhanced the efficacy of both active ingredients in susceptible *An. gambiae* and *An. coluzzii* populations (Figure 2). As noted in resistant mosquitoes, PBO synergized the activity of acetamiprid more effectively than that of clothianidin. Within 72 h of exposure to acetamiprid, mortality rose from 32.0 ± 1.6% without PBO to 94.0 ± 2.0 % with the synergist (Fisher’s exact test, p < 0.001, OR = 32.5, 95% CI = 12.6-101) in *An. gambiae* Kisumu. Mean mortality against acetamiprid at 72 h also tripled with pre-exposure to PBO both in the colonized strain *An. coluzzii* Ngousso and in field-collected *An. coluzzii* (95.0 ± 3.0 % and 45 ± 5.3 %, respectively). PBO did not significantly improve the efficacy of clothianidin against *An. gambiae* Kisumu (46.0 ± 12.8 % mortality without PBO vs 50.0 ± 17.0 % with PBO, Fisher’s exact test, p = 0.671, OR = 1.17, 95% CI = 0.65-2.12) nor in *An. coluzzii* Ngousso (41.0 ± 13.9 % vs 56.0 ± 8.6 %, p = 0.05, OR = 1.83, 95% CI = 1.00-3.34). However, mortality against clothianidin rose significantly with pre-exposure to PBO in field-collected *An. coluzzii* (18.0 ± 5.0 % vs 51.0 ± 16.1%, Fisher’s exact test, p < 0.001, OR = 4.70, 95% CI = 2.39-9.58).

**Figure 2:**
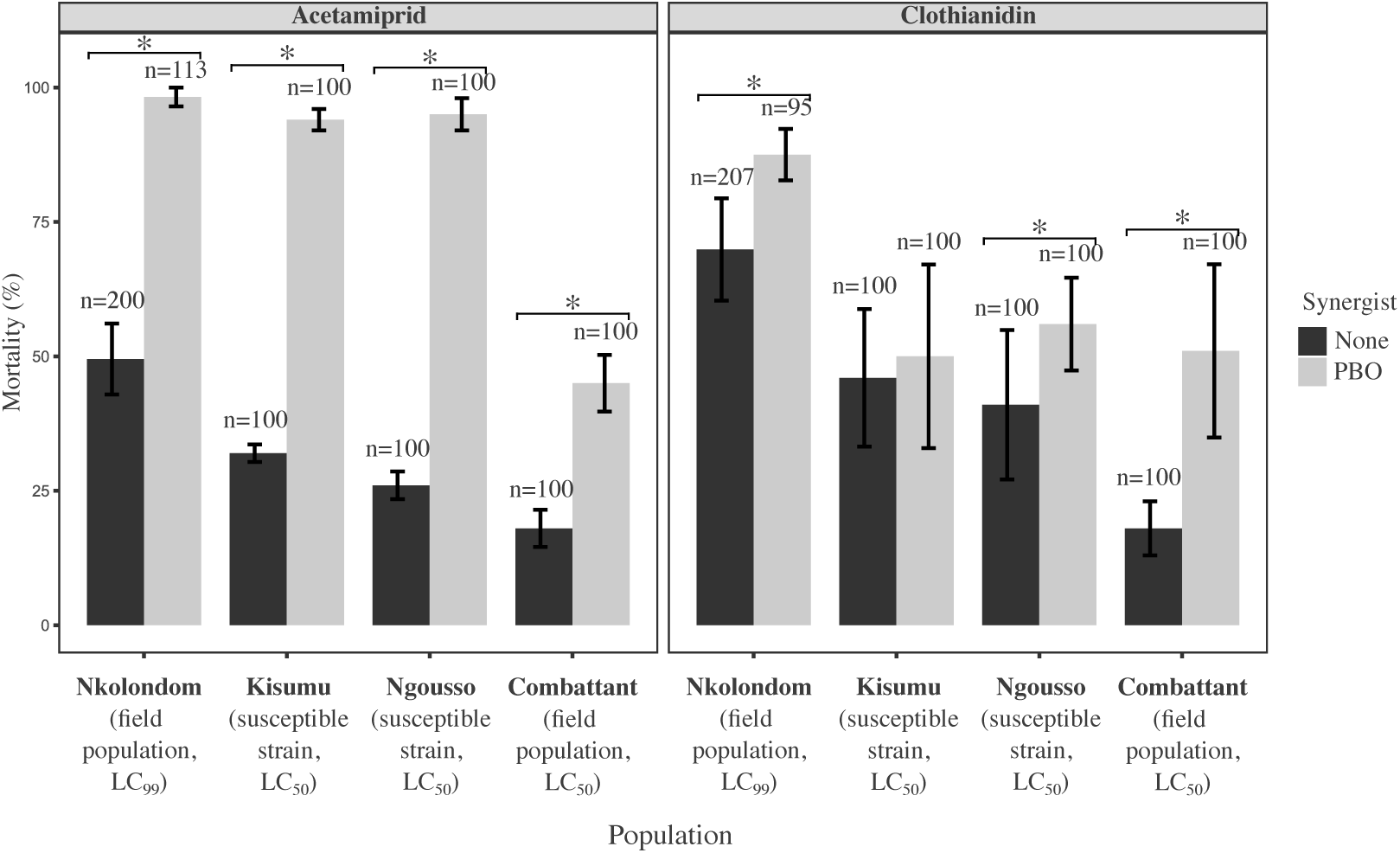
Synergistic effect of PBO on the activity of clothianidin and acetamiprid in *An. gambiae* and *An. coluzzii* adults. Mortality values were compared between tests exposing adult mosquitoes to the active ingredient alone (LC_99_ or LC_50_) and bioassays involving pre-exposure to PBO. Error bars represent the standard error of the mean. Asterisk denotes a significant increase in mortality due to PBO (Fisher’s exact test, p < 0.05).

### 3-3 Allele frequencies of nucleotide substitutions in Agβ1 and Agα6 subunits are correlated with sensitivity to neonicotinoid

Nucleotide substitutions and amino acid changes were detected in exons of Agβ1 and Agα6, and their allele frequencies compared between populations displaying a gradient of sensitivity to neonicotinoids. Notably, the frequency of an aspartic acid/glutamic acid substitution (D202E) in loop F of Agα6 varied according to the gradient of susceptibility described between *An. coluzzii* and *An. gambiae* (Figure 3). An aspartic acid was fixed in the most susceptible population, *An. coluzzii* Ngousso (n = 18). The frequency of this allele was also very high (83.33%) in susceptible *An. coluzzii* field populations (n = 30) and very low in resistant *An. gambiae* from Lobo and Nkolondom (13.51%) (n = 37) (Fisher’s exact test, p < 0.001, OR = 29.5, 95% CI = 7.19-154.7) as well as in individuals alive 72 h post-exposure to clothianidin or to acetamiprid (11.11%) (n = 36, ratio 1:1) (p < 0.001, OR = 36.4, 95% CI = 8.33-212.9). Still on Agα6, an isoleucine/methionine change (I198M) was detected on exon 7, but the association between its allele frequency and sensitivity was less striking (Figure 3). Precisely, although the frequency of isoleucine was 100% in *An. coluzzii* Ngousso suggesting that this allele increases sensitivity, the proportion of individuals bearing a methionine did not significantly vary between the susceptible strain *An. gambiae* Kisumu (38.10%) (n = 20) and resistant field-collected mosquitoes (51.43%) (Fisher’s exact test, p = 0.41, OR = 0.59, 95% CI = 0.17-1.99). A synonymous substitution whose frequencies increased gradually from susceptible to resistant populations was detected on a serine at position 201 of exon 7 (Figure 3). The adenine nucleotide was fixed in *An. coluzzii* Ngousso and very high (76.67%) in field-collected *An. coluzzii* indicating a correlation with susceptibility. Consistent with this hypothesis, the frequency of the adenine allele dropped to 24.32% and 27.78% respectively in non-exposed *An. gambiae* field-collected females and in individuals alive 72 h post-exposure to clothianidin or to acetamiprid. Another synonymous substitution segregating between *An. gambiae* and *An. coluzzii* was present on a leucine at position 218 of exon 7. A guanine was fixed in all *An. coluzzii* populations and occurred at less than 10% in field-collected *An. gambiae* mosquitoes suggesting a correlation between this mutation and susceptibility to neonicotinoids. Only two mutations that were synonymous substitutions were identified on three exons sequenced for the β1 subunit (Figure 4). On a valine at position 52, a cytosine was fixed in field-collected and laboratory *An. coluzzii*. Meanwhile, the same allele occurred at less than 15% in resistant *An. gambiae* field populations. An adenine/cytosine substitution was also found on another valine at position 100. Although the mutation was synonymous, a correlation with the gradient of sensitivity to neonicotinoids was obvious. The adenine allele was fixed in susceptible populations (*An. coluzzii* and *An. gambiae* Kisumu) whereas *An. gambiae* field populations were polymorphic with some heterozygous individuals.

**Figure 3:**
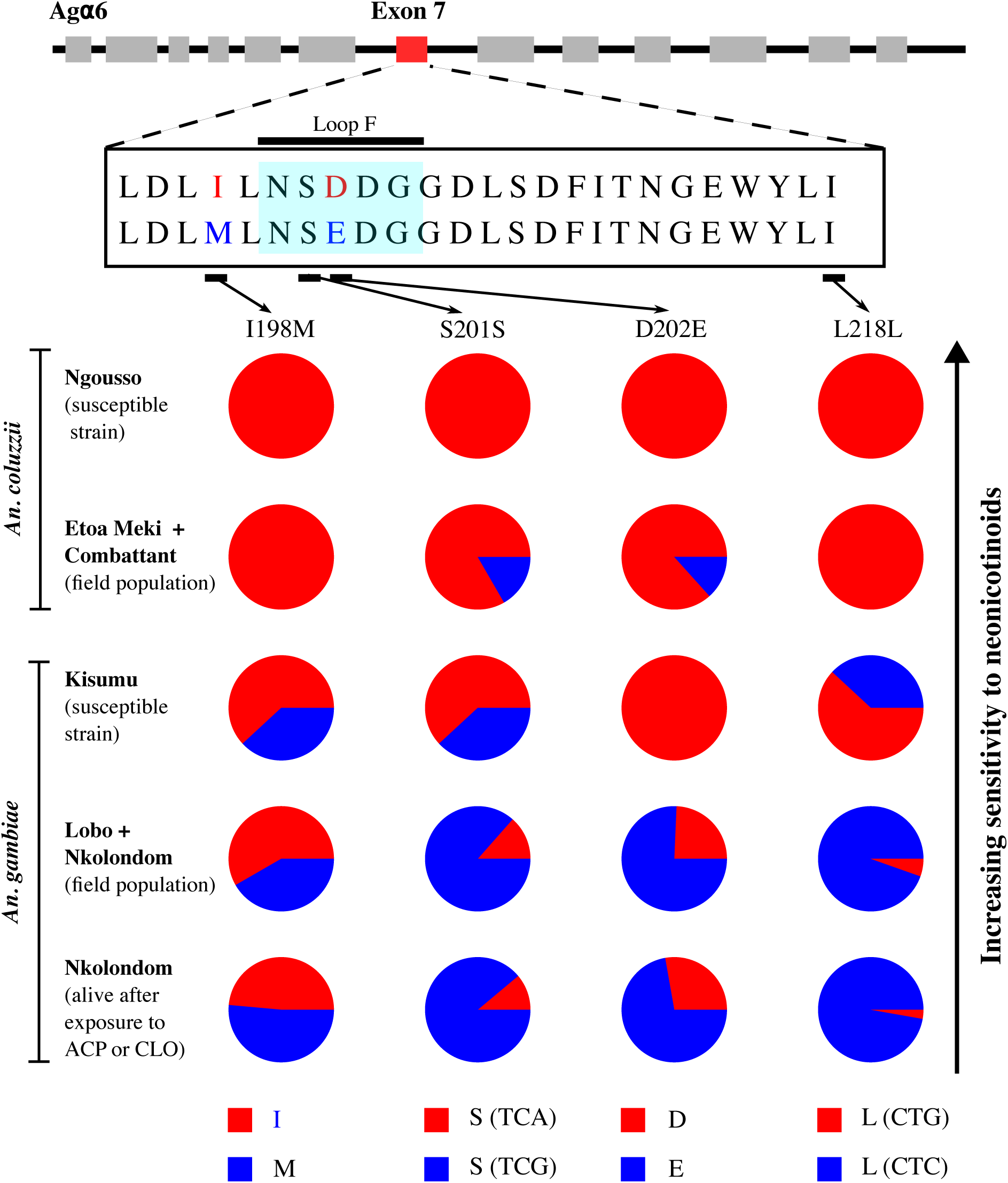
Distribution of allele frequencies of point mutations detected within and around Agα6 loop F. Two amino acid changes and two synonymous mutations are segregating between *An. gambiae* and *An. coluzzii*. A gradual rise in frequency from susceptible insectary colonies to resistant field populations indicates a correlation between the mutations and sensitivity to neonicotinoids. ACP = acetamiprid and CLO = clothianidin.

**Figure 4:**
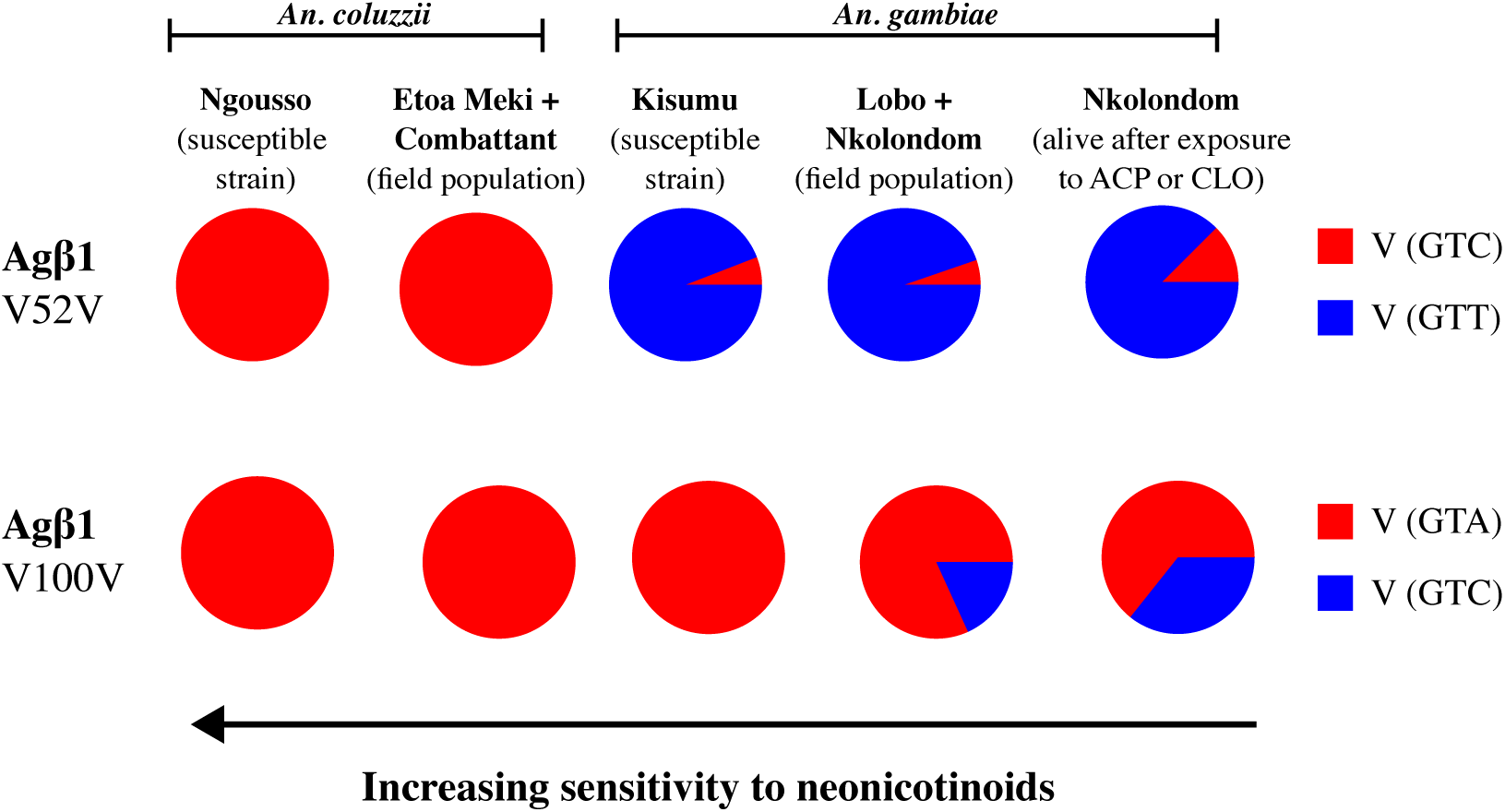
Allele frequencies of two synonymous mutations detected on Agβ1. A clinal distribution of frequency between *An. gambiae* and *An. coluzzii* suggests a correlation between the point mutations and sensitivity to neonicotinoid. ACP = acetamiprid and CLO = clothianidin.

### 3-4 The level of expression of nAChRs is reduced in *An. gambiae*

mRNA expression levels of eight subunits were compared between colonized susceptible strains and field-collected *An. gambiae* and *An. coluzzii* populations. The baseline level of expression of β2 was not significantly different between *An. gambiae* Kisumu and field-collected conspecific mosquitoes from Lobo (Welch t-test, p = 0.70, t = -0.42, df = 2.47) and from Nkolondom (p = 0.60, t = -0.60, df = 2.57) (Figure 5). Similarly, there was no significant difference in the level of expression of α9-1 between *An. gambiae* Kisumu and female adults from Lobo (Welch t-test, p = 0.27, t = -1.31, df = 3.45) and from Nkolondom (p = 0.71, t = -0.40, df = 3.39). On the contrary, the mRNA levels of β1, α3 and α7 decreased dramatically, on average 23.27 (p < 0.05, t = -3.53, df = 3.68), 17.50 (p < 0.05, t = -4.16, df = 3.66), and 15.80-fold (p < 0.05, t = -3.80, df = 4.0), respectively, in each *An. gambiae* field populations compared to Kisumu. The levels of expression were stable among field populations as no significant fold change was detected between females from Nkolondom and those from Lobo (Welch t-test, p > 0.05, except β1 with p = 0.04, t = 3.51, df = 2.99). Apart from a moderate decrease in mRNA concentration of β2 (2.67-fold change, p < 0.05, t = -3.47, df = 3.99) and α9-1 (5.28, p = 0.03, t = -5.20, df = 2.22) in Etoa Meki compared to the susceptible strain Ngousso, there was no significant variation in the level of expression of nAChRs across all pairwise comparisons between *An. coluzzii* populations (Figure 5).

**Figure 5:**
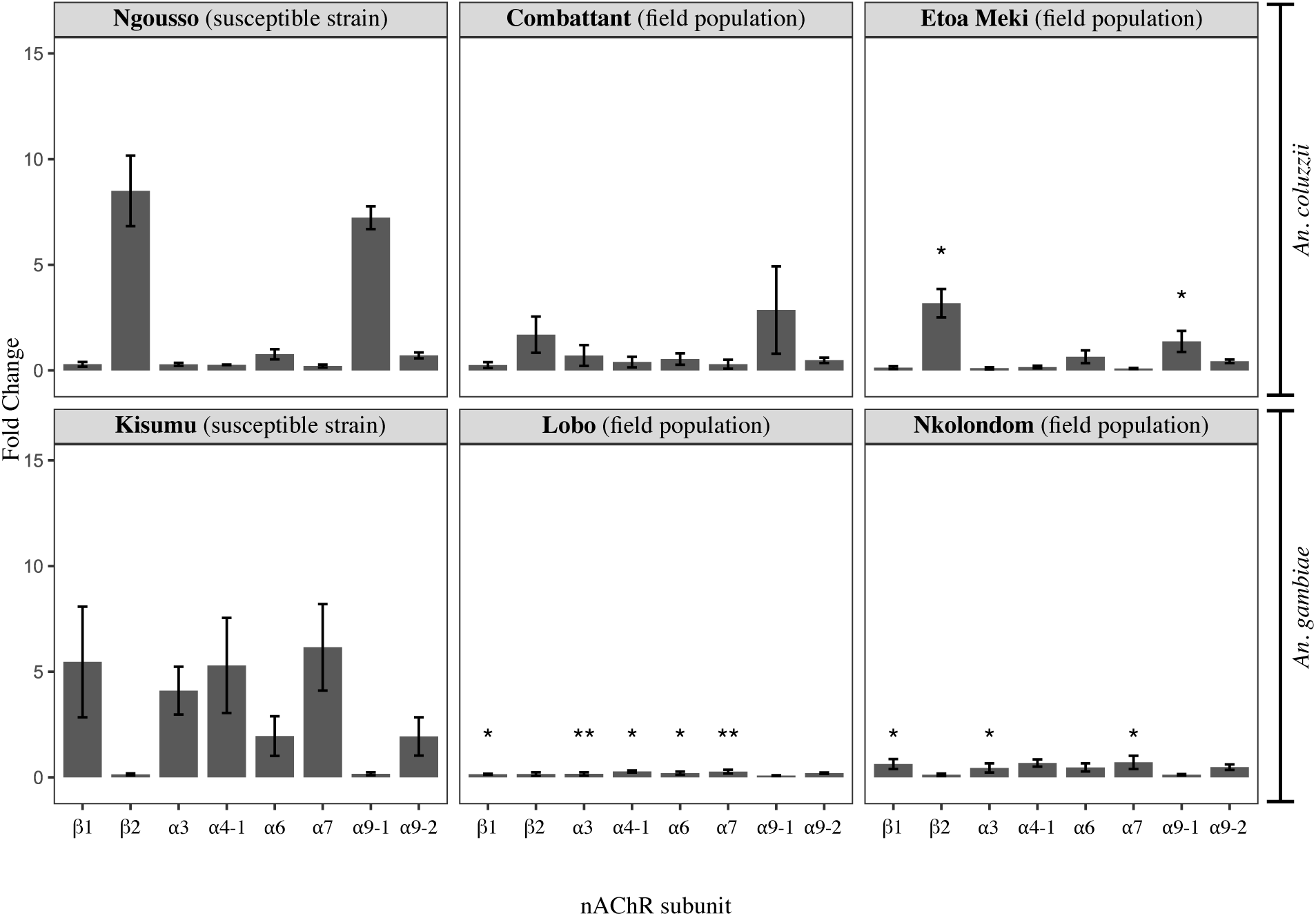
Relative mRNA expression level of nAChR subunits in *An. gambiae* and *An. coluzzii* adults displaying a gradient of susceptibility to clothianidin and acetamiprid. Error bars represent the standard error of the mean. The level of significance is indicated by *(p < 0.05-0.01) and **(p < 0.01-0.001).

## Discussion

Here we explored the mechanisms underlying the differential susceptibility to neonicotinoids observed in two *Anopheles* cryptic species. Results of adult bioassays are consistent with previous studies indicating that *An. coluzzii* populations from Yaounde are more susceptible to acetamiprid and clothianidin than is *An. gambiae* [27–29]. Contrary to *An. coluzzii* which is adapted to urbanization in this area, *An. gambiae* occurs in rural settings where larval populations are exposed to neonicotinoid residues which can drive resistance selection [28]. Dozens of formulations of acetamiprid, imidacloprid, thiacloprid and thiamethoxam are registered and freely used for crop protection in Cameroon [64,65]. Previous investigations also highlighted the primary role of cytochrome P 450s (CYPs) in the development of neonicotinoid resistance in *An. gambiae* [27,29] consistent with findings from studies involving diverse insect pests [47,48,50,54,55,66,67]. The novelty in the present study is the demonstration that metabolic detoxification based on CYPs is a pervasive adaptive mechanism that exists even in more sensitive populations such as susceptible laboratory strains. It also clearly appeared that, based on the synergistic effect of PBO, resistance to acetamiprid was more reliant on CYPs than was adaptation to clothianidin. The efficacy of CYP-mediated neonicotinoid metabolism varies depending on species and active ingredients. Notably, acetamiprid and clothianidin belong to two subclasses of neonicotinoids with different chemical characteristics, which could explain why acetamiprid is more effectively metabolized than clothianidin in *Anopheles* [24]. Studies across diverse taxa have provide evidence that contrary to acetamiprid, clothianidin is only partially metabolized by CYPs [67,68].

Wild *An. gambiae* mosquitoes exhibit reduced susceptibility to clothianidin, an active ingredient which is not effectively detoxified by CYPs suggesting that anopheline populations can evolve more complex resistance mechanisms including target-site insensitivity. At least nine nicotinic acetylcholine receptor subunits (nAChRs) have been identified in *An. gambiae*, but their affinity to different neonicotinoid insecticides as well as their role in resistance development have yet to be investigated [69]. This study revealed that difference in sensitivity to neonicotinoids between two sibling species is likely due in part to nucleotide substitutions and amino acid changes that potentially affect the interaction between nAChRs and active ingredients. Although nAChRs are relatively well conserved across taxa, mutations involved in resistance to neonicotinoids, to sulfoxaflor or to spinosad have been detected in at least four subunits in insect pests [23]. We have identified four synonymous substitutions and two amino acid changes exhibiting clines of allelic frequencies in wild *Anopheles* populations within the β1 and α6 subunits. Four amino acid substitutions whose role in reducing sensitivity to neonicotinoids has been established through binding affinity tests, electrophysiological analyses and functional studies are known on the β1 subunit in agricultural pests [35,36,40,41,70]. In the present study, none of the four amino acid substitutions was detected in *Anopheles* mosquitoes from Yaounde. Instead, two synonymous mutations whose allele frequencies were correlated with a gradient of susceptibility were observed. It remains to be established whether these nucleotide substitutions play a functional role in susceptibility to neonicotinoids, but synonymous mutations might be a tradeoff enabling a subtle reduction in sensitivity of the target site while minimizing the fitness cost that could be associated with a more profound change such as an amino acid substitution [45].

However, on the Agα6 subunit, two amino acid changes were detected in field-collected mosquitoes and their allele frequencies were also associated with a gradient of susceptibility between *An. gambiae* and *An. coluzzii*. The G275E substitution known to increase resistance to sipinosad was not present in *Anopheles* mosquitoes, but one of the amino acid changes found on Agα6 (D202E) fell in loop F. The tridimensional structure of nAChR subunits harbor six loops (A to F) known to modulate the binding affinity of ligands; loop A to D acting on α subunits and loop D to F affecting non-α subunits. This study provides an instance of amino acid change with potential implications for the binding of neonicotinoids occurring on loop F of an α subunit. Although the role of the D202E mutation remains unknown, these findings suggest that loop F may play an important role in binding in both α and non-α subunits in some insect species [23].

Agα6 also contained two synonymous mutations showing a correlation with sensitivity to neonicotinoids in *Anopheles* mosquitoes. Overall, allele frequencies of four synonymous substitutions present on Agβ1 and Agα6 are segregating between *An. coluzzii* and *An. gambiae* and are associated with resistance to acetamiprid and clothianidin. Previous studies screening molecular variations across both subunits did not report the presence of synonymous mutations in agricultural pests [39,42,61,71]. One possible explanation of the discrepancy between other investigations and the current findings is that most studies are based on insecticide resistance selection using colonized insect populations. This approach can lead to an underestimation of polymorphism across nAChRs. Our study describes an instance of field-evolved resistance that may trigger more complex mechanisms including synonymous substitutions. The role of synonymous mutations in protein evolution remains unclear, but such changes are known to alter phenotype by affecting transcription and translation, RNA splicing and transport, or codon usage [72].

As neonicotinoid insecticides differentially target nAChR subunits, selective regulation of expression levels is another known mechanism that affects sensitivity in insects. Studies have revealed remarkable examples of reduced expression of one or several nAChR subunits correlated with resistance development. Notably, It has been shown that knockout of the α6 subunit increases resistance to spinosad while deletion of α1 confers resistance to dinotefuran in *Drosophila melanogaster* [45,46]. It was also found that the α8 subunit was substantially downregulated in an imidacloprid-resistant population of the brown planthopper, *Nilaparvata lugens* compared to susceptible individuals [43]. The present study revealed that field-evolved resistance to clothianidin and acetamiprid is associated with reduced expression of the β1, α3, and α7 and subunits in *An. gambiae*. Thiamethoxam resistance in *Aphis gossypii* also involves a reduction in mRNA expression levels of α1, α4-1, α4-2 and α5 and α7.

Evidence from field-evolved resistance to neonicotinoids in *Anopheles* mosquitoes corroborates findings from several studies indicating that when metabolic detoxification is insufficient, insect pests can use a combination of molecular variants and transcriptional regulations to alter the sensitivity of the target site [23]. This study explored variations across subsets of two out of at least nine nAChR subunits present in *Anopheles* mosquitoes. A more comprehensive analysis of mutations within subunits and their association with binding affinity and neonicotinoid sensitivity in the major vectors of *Plasmodium* in Africa will be the focus of further investigation.

In summary, the findings of this study provide evidence that wild *An. gambiae* populations are evolving cross-resistance to several neonicotinoids facilitated by metabolic resistance and a reduction in sensitivity of the target site. Exploring the selective pressures and molecular mechanisms underlying susceptibility in *Anopheles* will provide the scientific basis for efficient application of neonicotinoids in vector control.

## Acknowledgements

We thank several helpers for their assistance in larval collection.

## Author contributions

Conceptualization: Colince Kamdem, Caroline Fouet, Matthew J Pinch. Formal analysis: Caroline Fouet, Fred A. Ashu, Colince Kamdem. Funding acquisition: Colince Kamdem. Investigation: Caroline Fouet, Fred A. Ashu, Marilene M. Ambadiang, Matthew J Pinch, Colince Kamdem. Methodology: Fred A. Ashu, Caroline Fouet, Marilene M. Ambadiang, Calmes Bouaka, Matthew J Pinch, Anthoni J. Bathronie, Cesar A. Hernandez, Desiree E. Rios. Project administration: Colince Kamdem. Resources: Veronique Penlap-Beng. Supervision: Colince Kamdem, Veronique Penlap-Beng. Writing–original draft: Caroline Fouet, Colince Kamdem. Writing–review & editing: Colince Kamdem

